# Comparative Pathogenesis of Different Phylogroup I Bat Lyssaviruses in a Standardized Mouse Model

**DOI:** 10.1101/2021.09.29.462306

**Authors:** Antonia Klein, Elisa Eggerbauer, Madlin Potratz, Luca M. Zaeck, Sten Calvelage, Stefan Finke, Thomas Müller, Conrad M. Freuling

## Abstract

A plethora of bat-associated lyssaviruses potentially capable of causing the fatal disease rabies are known today. Transmitted via infectious saliva, occasionally-reported spillover infections from bats to other mammals demonstrate the permeability of the species-barrier and highlight the zoonotic potential of bat-related lyssaviruses. However, it is still unknown whether and, if so, to what extent, viruses from different lyssavirus species vary in their pathogenic potential. In order to characterize and systematically compare a broader group of lyssavirus isolates for their viral replication kinetics, pathogenicity, and virus release through saliva-associated virus shedding, we used a mouse infection model comprising a low (10^2^ TCID_50_) and a high (10^5^ TCID_50_) inoculation dose as well as three different inoculation routes (intramuscular, intranasal, intracranial). Clinical sings, incubation periods, and survival were investigated. Based on the latter two parameters, a novel pathogenicity matrix was introduced to classify lyssavirus isolates. Using a total of 13 isolates from ten different virus species, this pathogenicity index varied within and between virus species. Interestingly, Irkut virus (IRKV) and Bokeloh bat lyssavirus (BBLV) obtained higher pathogenicity scores (1.14 for IRKV and 1.06 for BBLV) compared to Rabies virus (RABV) isolates ranging between 0.19 and 0.85. Also, clinical signs differed significantly between RABV and other bat lyssaviruses. Altogether, our findings suggest a high diversity among lyssavirus isolates concerning survival, incubation period, and clinical signs. Virus shedding significantly differed between RABVs and other lyssaviruses. Our results demonstrated that active shedding of infectious virus was exclusively associated with two RABV isolates only (92 % for RABV-DogA and 67 % for RABV-Insectbat), thus providing a potential explanation as to why sustained spillovers are solely attributed to RABVs. Interestingly, high-resolution imaging of a selected panel of brain samples from bat-associated lyssaviruses demonstrated a significantly increased percentage of infected astrocytes in mice inoculated with IRKV (10.03 %; SD±7.39) compared to RABV-Vampbat (2.23 %; SD±2.4), and BBLV (0.78 %; SD±1.51), while only individual infected cells were identified in mice infected with Duvenhage virus (DUVV). These results corroborate previous studies on RABV that suggest a role of astrocyte infection in the pathogenicity of lyssaviruses.

**Author Summary:** Globally, there are at present 17 different officially recognized lyssavirus species posing a potential threat for human and animal health. Bats have been identified as carriers for the vast majority of those zoonotic viruses, which cause the fatal disease rabies and are transmitted through infectious saliva. The occurrence of sporadic spillover events where lyssaviruses are spread from bats to other mammalian species highlights the importance of studying pathogenicity and virus shedding in regard to a potentially sustained onward cross-species transmission. Therefore, as part of this study, we compared 13 different isolates from ten lyssavirus species in a standardized mouse infection model, focusing on clinical signs, incubation periods, and survival. Based on the latter two, a novel pathogenicity index to classify different lyssavirus species was established. This pathogenicity index varied within and between different lyssavirus species and revealed a higher ranking of other bat-related lyssaviruses in comparison to the tested Rabies virus (RABV) isolates. Altogether, our results demonstrate a high diversity among the investigated isolates concerning pathogenicity and clinical picture. Furthermore, we comparatively analyzed virus shedding via saliva and while there was no indication towards a reduced pathogenicity of bat-associated lyssaviruses as opposed to RABV, shedding was increased in RABV isolates. Additionally, we investigated neuronal cell tropism and revealed that bat lyssaviruses are not only capable of infecting neurons but also astrocytes.

## 1. Introduction

The *Lyssavirus* genus of the family *Rhabdoviridae* within the order *Mononegavirales* comprises highly neurotropic, single negative-strand RNA viruses [1], which are capable of causing rabies, an acute and invariably fatal viral encephalitis [2]. At present, 17 lyssavirus species are recognized as separate taxonomic entities [1]. Based on antigenic divergence and phylogenetic relationships, lyssavirus species can be grouped into phylogroups [3]. Phylogroup I includes the prototypical Rabies virus (RABV), Aravan virus (ARAV), Australian bat lyssavirus (ABLV), Bokeloh bat lyssavirus (BBLV), Duvenhage virus (DUVV), European bat lyssavirus 1 (EBLV-1), European bat lyssavirus 2 (EBLV-2), Gannoruwa bat lyssavirus (GBLV), Irkut lyssavirus (IRKV), Khujand lyssavirus (KHUV), and Taiwan Bat Lyssavirus (TWBLV), while Lagos bat lyssavirus (LBV), Mokola lyssavirus (MOKV), and Shimoni bat lyssavirus (SHIBV) are members of phylogroup II. Based on phylogenetic distance, the most genetically divergent lyssaviruses, including Ikoma virus (IKOV), Lleida bat lyssavirus (LLEBV), and West Caucasian bat lyssavirus (WCBV), have been tentatively classified within a dispersed phylogroup III [4]. Two new lyssaviruses found in Europe, the Kotalahti bat lyssavirus (KBLV) [5], and Africa, the Matlo bat lyssavirus (MBLV) [6], are not yet approved as new virus species. While almost all lyssavirus species are strongly associated with chiropteran hosts [4], RABV is the only lyssavirus maintained in many different species of mesocarnivores around the world. Exceptions include the circulation of RABV in multiple species of bats in the New World and the reported role of a small primate, the marmoset, as an RABV reservoir in Brazil [2,7]. Except for bat-associated RABV in the Americas and ABLV, all other bat lyssaviruses seem to be restricted to one reservoir host species they have been steadily co-evolving with over time [8].

For RABV, transmission, particularly from vampire bats, to other non-bat mammals is common in the Americas [9]. Nonetheless, sporadic spillover infections of other bat lyssaviruses to non-bat mammal species, including humans, emphasize the threat these viruses pose for both human and animal health [10,11]. In contrast to phylogroup I lyssaviruses, failure of protection against the more divergent phylogroup II and III lyssaviruses has been demonstrated for all commercially available vaccines [3,12– 14]. While antigenicity and phylogeny of lyssaviruses have been well studied, the comparative pathogenicity of bat lyssavirus isolates in different species is of scientific interest, particularly against the background of a highly variable pathogenicity between phylogroups observed in mice [15]. Partly, pathogenicity of bat lyssaviruses was tested in ferrets [16], foxes [17–19], raccoon dogs [18,20–22], cats [18,21–28], dogs [18,21,22,24,25,29–32], and skunks [18,21,22,33]. In most of these studies and also when mice were used, either only a limited number of bat lyssavirus species were compared or different viral variants of one particular lyssavirus species have been studied [34,34–40]. Varying experimental designs and conditions often prevent a reliable and broader comparative assessment of the pathogenicity of bat lyssaviruses from these studies. However, in many of the aforementioned sstudies, the results regarding limited pathogenicity of bat lyssaviruses in non-bat mammals seem to contradict reported bat lyssavirus-borne human fatalities [10].

Moreover, it has not yet been clarified why cross-species transmissions to either humans or animals are more often seen with bat-associated RABVs [8] compared to other bat lyssaviruses for which such events seem to be relatively rare [10]. While the underlying mechanisms of cross-species transmissions are not yet completely understood, virus shedding is assumed to be a key factor, particularly for sustained spillovers [41]. Therefore, it is of importance to understand whether potential spillover hosts are shedding virus and can subsequently transmit it to conspecifics or other mammals, including humans. Such assessment of the potential impact of bat-associated lyssaviruses on public health is particularly challenging but of great importance. To this end, we compared the pathogenicity of 13 phylogroup I lyssaviruses in a standardized mouse model using different inoculation doses and routes. Furthermore, shedding of virus in saliva of infected mice was measured in order to assess the likelihood of onward transmission. Based on obtained pathogenicity data, we further developed a pathogenicity index for comparison and classification of lyssavirus-induced pathogenicity. Since it was shown that the degree of central nervous system resident astrocyte infection differed in RABV field strains compared to laboratory-adapted fixed virus strains, potentially affecting their pathogenicity [42], the cell tropism of a selected lyssavirus panel from diseased mice was analyzed. Even though the restricted number of lyssaviruses analyzed for astrocyte infection hinders a full comparison, the results support previous studies of RABV on the association of astrocyte tropism and pathogenicity.

## 2. Material and Methods

### 2.1 Ethics statement

The experimental work in mice strictly followed the European guidelines on animal welfare and care according to the authority of the Federation of European Laboratory Animal Science Associations (FELASA) [43]. Animal experiments were evaluated, reviewed, and approved by the animal welfare committee (Landesamt für Landwirtschaft, Lebensmittelsicherheit und Fischerei Mecklenburg-Vorpommern, LALLF M-V/TSD 7221.3-2.1-002/11; M-V/TSD/7221.3-2-001/18) and supervised by the commissioner for animal welfare at the Friedrich-Loeffler-Institut (FLI) representing the Institutional Animal Care and Use Committee (IACUC).

### 2.2 Viruses and cells

A total of 13 virus isolates representing ten different phylogroup I lyssaviruses originating from Europe (EBLV-1, EBLV-2, BBLV), Asia (ARAV, KHUV, IRKV, GBLV), Africa (DUVV), Australia (ABLV), and the Americas (RABV) were included in this study. Regarding the latter, two bat-related RABV isolates, one from an insectivorous (RABV-Insectbat) and one from a hematophagous bat (RABV-Vampbat) (Table 1), were selected. For EBLV-1, DUVV, and ABLV, isolates from human cases were used. The Asian bat lyssaviruses ARAV, IRKV, GBLV and KHUV were kindly provided by the Animal Plant and Health Agency (APHA), Weybridge, United Kingdom through the European Virus Archive global (EVAg). All other isolates originated from the virus archive of the FLI, Riems, Germany. For comparison, two representatives of classical non-bat RABVs, one being an isolate from a dog from Azerbaijan (RABV-DogA) and the other being a wildlife variant isolated from a raccoon from North America (RABV-Raccoon), both of which had been used in previous infection studies in raccoons [20], were included. Cell lines used in this study were obtained from the Collection of Cell Lines in Veterinary Medicine (CCLV; FLI, Riems, Germany). Mouse neuroblastoma cells (Na 42/13, CCLV-RIE 0229) maintained in a mixture of equal volumes of Eagle MEM (Hanks’ balanced salts solution) and MEM (Earle’s balanced salts solution) medium supplemented with 10 % fetal bovine serum and penicillin/streptomycin (100 U/ml and 100 µg/ml, respectively) were used for propagation of virus stocks, titration, viral replication kinetics, and virus isolation.

**Table 1:** Viruses used in the study, including details of their year of isolation, the respective host, and origin.

| Lab-ID | Name | Virus species | Year | Host | Origin | Accession Number |
| --- | --- | --- | --- | --- | --- | --- |
| 5989 | RABV-DogA | RABV | 2002 | dog (*Canis lupus familiaris*) | Azerbaijan | LN879480 |
| 13205 | RABV-Raccoon | RABV | 1981 | raccoon (*Procyon lotor*) | United States | MN862283 |
| 34886 | RABV-Vampbat | RABV | 1973 | vampire bat (*Desmodus rotundus*) | Latin America | PRJEB46947 |
| 13240 | RABV-Insectbat | RABV | 1986 | big brown bat (*Eptesicus fuscus*) | Canada | PRJEB46947 |
| 13027 | EBLV-1 | EBLV-1 | 1982 | human (Yuli virus) | Russia | LT839613 |
| 16618 | EBLV-2 | EBLV-2 | 2007 | daubenton's bat (*Myotis daubentonii*) | Germany | KY688138 |
| 29008<sup>#</sup> | BBLV | BBLV | 2010 | natterer's bat (*Myotis Nattererii*) | Germany | KF245925 |
| 46579 | ARAV | ARAV | 1991 | lesser mouse-eared bat (*Myotis blythii*) | Kyrgyzstan | EF614259 |
| 46580 | KHUV | KHUV | 2001 | whiskered bat (*Myotis mystacinus*) | Tajikistan | NC025385 |
| 46582 | IRKV | IRKV | 2002 | greater tube-nosed bat (*Murina leucogaster*) | Russia | NC020809 |
| 39663 | GBLV | GBLV | 2016 | indian flying fox (*Pteropus medius*) | Sri Lanka | KU244267 |
| 12862 | DUVV | DUVV | 1971 | human | South Africa | EU293119 |
| 13849 | ABLV | ABLV | 1986 | human | Australia | AF418014 |

<sup>#</sup>A recombinant virus was used.

### 2.3 Full genome sequencing

All viruses taken from archived samples were subjected to full genome sequencing essentially as described before [44]. Briefly, RNA was automatically extracted on a KingFisher Flex platform (Thermo Fisher Scientific, Waltham, MA, USA) using the RNAdvance Tissue Kit (Beckmann Coulter, Brea, CA, USA). Double stranded cDNA was generated from 350 ng total RNA using the SuperScript™ IV First-Strand cDNA Synthesis System (Invitrogen/Thermo Fisher Scientific, Waltham, MA, USA) and the NEBNext® Ultra™ II Non-Directional RNA Second Strand Synthesis Module (New England Biolabs, Ipswich, MA, USA). After conversion into cDNA, fragmentation was achieved by ultrasonication on a Covaris M220 (Covaris, Brighton, UK). Subsequently, Ion Torrent-specific sequencing libraries were generated using the GeneRead L Core Kit (Qiagen, Hilden, Germany) together with IonXpress barcode adaptors (Thermo Fisher Scientific). After quantification (QIAseq Library Quant Assay Kit, Qiagen) and quality control (2100 Bioanalyzer, High sensitivity DNA Kit, Agilent Technologies, Santa Clara, CA, USA) of the libraries, sequencing was performed on an Ion Torrent S5XL instrument utilizing Ion 530 chips and reagents according to the manufacturer’s instructions.

### 2.4 Viral propagation and replication kinetics

In order to generate virus stocks for inoculation, Na 42/13 cells were infected at a multiplicity of infection (MOI) of 0.001, and incubated at 37°C and 5 % CO_2._ Depending on the viral strain, supernatants were harvested 72 to 168 hours post-infection (hpi) when 100 % of the cell monolayer was infected. Infection was assessed using a control dish stained with a fluorescein isothiocyanate (FITC)-conjugated monoclonal antibody mix (SIFIN, Germany/Fujirebio, Belgium). For growth curves, Na 42/13 cells were infected at an MOI of 0.001. Cell culture supernatants were harvested at 0, 16, 24, 48, 72, 96, and 168 hpi. The virus titers (tissue culture infective dose 50 - TCID_50_) were determined by endpoint titration of three technical replicates and subsequent calculation by the Spearman-Karber method [45].

### 2.5 Animal experiments

For the experimental studies, three-to four-week-old Balb/c mice from a commercial breeder (Charles River, Germany) were used. Animals were randomly assigned to groups and housed in individual, labelled cages with water and food provided ad libitum. Six mice each were inoculated intramuscularly (i.m.) in the femoral muscle using a high (10^5^ TCID_50_/30 µl) and a low (10^2^ TCID_50_/30 µl) viral dose. Additionally, one group of six and one of three mice was inoculated intranasally (i.n.) with 10^2^ TCID_50_/10 µl and intracranially (i.c.) with 10^2^ TCID_50_/30 µl, respectively. Mock-infected control groups for each administration route were inoculated with 10 µl or 30 µl of cell culture medium respectively. Animals were monitored for 21 days post-infection (dpi). Body weight and clinical signs were recorded daily for each animal using clinical scores (S1 Fig). If animals showed more than one clinical sign at a given time point, the most prominent one dominating the physical condition was recorded. Once mice developed clinical signs, they were checked twice a day. Animals were humanely euthanized at the humane endpoint or after 21 days by cervical dislocation under anesthesia with isoflurane (Isofluran CP, CP Pharma, Germany). Immediately before euthanasia, oropharyngeal swabs were taken of all mice that succumbed to infection using dry sterile cotton swabs (Nerbe plus GmbH & Co. KG, Germany), which were placed into 1500 μl of cell culture medium supplemented with penicillin/streptomycin as described above. Upon euthanasia, salivary glands and brain samples of all animals were taken and stored at -80° C until further processing.

### 2.6 Diagnostic assays

Presence of lyssavirus antigen in heat-fixed brain tissue samples was detected by direct fluorescence antibody test (DFA) using FITC-conjugated monoclonal antibodies (SIFIN, Germany and Fujirebio, Belgium) as well as defined positive and negative controls [46].

Brain, salivary gland and oropharyngeal swab samples were used to detect lyssaviral RNA. Briefly, organ samples were homogenized in 1000 µl cell culture medium using a TissueLyser (Qiagen, Germany) with a 3 mm steal bead. Homogenates as well as oral swabs were centrifuged at 3750 x g for 10 minutes. Viral RNA was then extracted from the supernatant (100 µl) using the NucleoMagVet kit (Macherey&Nagel, Germany) according to the manufacturer’s instructions in a KingFisher/BioSprint 96 magnetic particle processor (Qiagen, Germany). Viral RNA was detected by an RT-qPCR targeting the N-gene using the R14-assay (RABV-, EBLV-1-, EBLV-2-, and BBLV) and the R13-assay (ABLV, DUVV) [47,48]. For GBLV, primers and probe were specifically designed (Table S1) and the protocol was run separately but with the same conditions as below. The PCR mastermix was prepared using the AgPath-ID One-step RT-PCR kit (Thermo Fisher Scientific, USA) in a volume of 10 µl including 0.5 µl of β-Actin-mix2-HEX as internal control and 2.5 µl of extracted RNA. The reaction was performed for 10 minutes at 45°C for reverse transcription and 10 minutes at 95°C for activation, followed by 42 cycles of 15 seconds at 95°C for denaturation, 20 seconds at 56°C for annealing and 30 seconds at 72°C for elongation. Fluorescence was measured during the annealing phase. RNA specific for ARAV, IRKV and KHUV was detected using the pan-lyssa realtime PCR targeting both the N- and L-gene [47,48]. Here, the RT-qPCR reaction was prepared using the OneStep RT-PCR kit (Qiagen, Germany), adjusted to a volume of 12.5 µl with an internal control mastermix, based on β-Actin, running in parallel. The reaction consisted of 10 minutes at 45°C for reverse transcription and 10 minutes at 95°C for activation, followed by 45 cycles of 15 seconds at 95°C for denaturation, 20 seconds at 56°C for annealing and 30 seconds at 72°C for elongation. All RT-qPCRs were performed on a BioRad CFX96 Real-Time System (Bio-Rad, USA).

RT-qPCR positive salivary glands and oral swab samples were subjected to virus isolation in cell culture using the rabies tissue-culture infection test (RTCIT) [49]. Briefly, either the respective swap sample or supernatant from the homogenized organ suspension prepared using cell culture media as described above and supplemented with 10 % fetal bovine serum and 200 U/ml und 200 µg/ml penicillin/streptomycin, was mixed with dextran-pretreated Na 42/13 cell suspension at an equal ratio. The mixture was then incubated at 37°C and 5 % CO_2_ for 30 minutes and centrifuged at 1250 x g for 10 minutes. The obtained cell pellets were resuspended in T25 cell culture flasks and incubated for three to four days at 37°C and 5 % CO_2_. A control-dish was fixed, stained with a commercial FITC-conjugated monoclonal antibody conjugate (SIFIN, Germany/Fujirebio, Belgium), washed and microscopically analyzed for the presence of virus. Three consecutive serial passages were used to confirm a negative result.

### 2.7 Antibodies for immunofluorescence imaging of solvent-clearing brain sections

To detect bat lyssavirus antigen in infected brains, a polyclonal rabbit serum against recombinant RABV P protein (P160-5, 1:3,000 in PTwH [0.2 % Tween 20 in PBS with 10 µg/ml heparin]) was used [50]. The following commercial primary antibodies were used: chicken anti-GFAP (Thermo Fisher, USA; #PA1-1004, RRID:AB_1074620, 1:1,500 in PTwH) and guinea pig anti-NeuN (Synaptic Systems, Germany; #266004, RRID:AB_2619988, 1:800 in PTwH). Donkey anti-rabbit Alexa Fluor® 568 (Thermo Fisher, USA; #A10042, RRID:AB_2534017) and donkey anti-guinea pig Alexa Fluor® 647 (Dianova, Germany; #706-605-148, RRID:AB_2340476) were used as secondary antibodies, each at a dilution of 1:500 in PTwH.

### 2.8 Immunostaining and clearing of brain tissue samples

Immunostaining and clearing protocols from previous reports [51–53] were modified and performed as described previously [54]. All incubation steps were performed on an orbital shaker. Briefly, the PFA-fixed brains were cut into 1 mm thick slices using a vibratome (Leica Biosystems, Germany, VT1200S). After bleaching with 5 % H_2_O_2_/PBS overnight at 4°C the samples were pre-permeabilized (0.2 % Triton X-100/PBS) twice for 3 h each at 37°C and then permeabilized (0.2 % Triton X-100/20 % DMSO/0.3 M glycine/PBS) for 2 days at 37° C. After subsequent blocking (0.2 % Triton X-100/10 % DMSO /6 % donkey serum/PBS) at 37°C for further 48 h, primary antibodies diluted in 3 % donkey serum/5 % DMSO/PTwH were added for a total of 7 days at 37° C. After 3.5 days, the antibody solution was renewed once. Subsequently, the samples were washed with PTwH four times with increasing intervals, leaving the final wash on overnight. Secondary antibodies were diluted in 3 % donkey serum/PTwH and incubation was performed as described for the primary antibodies. After further washing with PTwH four times, leaving the final wash on overnight, the samples were dehydrated in a graded ethanol series (30 %, 50 %, 70 % in aqua ad iniectabilia [pH 9-9,5] and twice 100 %; each for ≥6 h) at 4° C. Subsequently, they were delipidated for 2 h in *n*-hexane at room temperature. Gradually replacing the *n*-hexane with ethyl cinnamate (ECi), they were then incubated in ECi until optically transparent. For confocal laser-scanning microscopy analysis, the cleared samples were embedded in 3D-printed imaging chambers as described before [55].

### 2.9 Confocal laser-scanning microscopy and image processing

Confocal z-stacks were acquired with a Leica DMI 6000 TCS SP5 confocal laser-scanning microscope equipped with a 40×/1.10 water immersion HC PL APO objective using the Leica Application Suite Advanced Fluorescence software (v2.7.3.9723). Fluorescence was acquired sequentially between lines with a pinhole diameter of 1 Airy unit and a z-step size of 0.5 µm.

The quantification of infected neurons and astrocytes in 1mm thick brain sections was done as according to previous description [42]. To this end, confocal image stacks were split into individual channels using Fiji, an ImageJ (v1.52h) distribution package [56]. After bleach correction (simple ratio, background intensity 5.0) brightness and contrast were adjusted for each channel. The 3D objects counter plugin was used to identify objects in each channel [57]. The resulting objects map was then overlaid with the RABV P channel to detect and count infected objects. Visualization was done using arivis vision4D (v3.4.0).

### 2.10 Statistical analysis

Statistical analyses were performed using Prism version 8 (GraphPad) with selection of the test as outlined in the results. P values of <0.05 were considered statistically significant.

### 2.11 Calculation of intramuscular pathogenicity index (IMPI)

For the calculation of the IMPI, we followed the example of the intracerebral pathogenicity index (ICPI) test for Newcastle disease virus [58] with the following modifications: The clinical observations of individual animals recorded every 12 to 24 hours over a period of up to 21 days were transferred to a daily rating scheme. Mice were scored as follows: 0 if healthy; 1 if sick, and 2 if dead. Dead individuals were scored as 2 at each of the remaining daily observations after death. For calculation of the IMPI, only animals from the i.m.-inoculated groups were included. The intramuscular pathogenicity index was then calculated based on the following formula: cumulative score for sick animals + cumulative score of dead animals / 126 (number of animals, i.e. 6 x days of observation, i.e. 21). The index is determined as the mean score per mouse over a 21-day-period, i.e. very pathogenic viruses showing high and less pathogenic ones showing lower indices. A minimum index value of 0 corresponds to absolutely apathogenic animals throughout the complete observation period while a maximum index value of 2 would be reached if all mice died at day 1 post-infection.

## 3. Results

### 3.1 In vitro replication kinetics

In mouse neuroblastoma cells (Na 42/13), the tested lyssaviruses replicated to maximum titers ranging from 10^5^ TCID_50_/ml (ABLV) to 10^7.75^ TCID_50_/ml (RABV-Vampbat) after 168 hours (Fig 1). RABV-Vampbat, IRKV, GBLV, RABV-DogA, and EBLV-1 exhibited titers around 10^7^ TCID_50_/ml and higher, while the titers for the rest of the isolates ranged between 10^5^ and 10^6^ TCID_50_/ml. IRKV and EBLV-1 showed the fastest replication with measurable titers obtained already after 16 and 24 hpi, while all other isolates yielded measurable titers either after 48 or 72 hpi. The RABV-Raccoon variant in particular exhibited a slow replication kinetic; replication started after 72 hpi but with comparably low titers even after 96 hpi (Fig 1).

**Fig 1:**
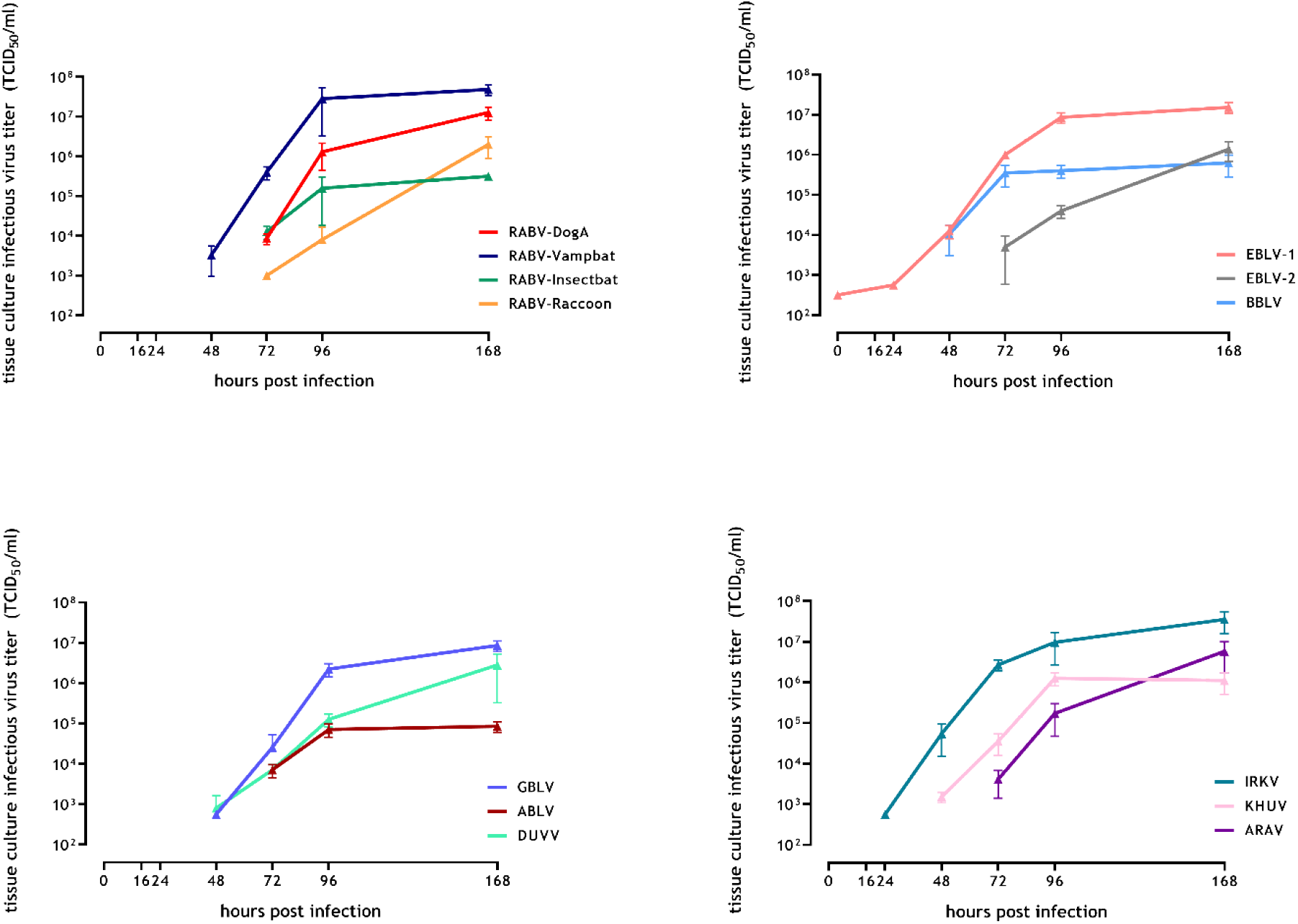
In vitro replication kinetics of lyssavirus isolates in Na 42/13 cells infected with an MOI of 0.001. The mean and standard errors of three replicates are indicated.

### 3.2 Survival rates

The survival rates of mice investigated according to the aforementioned experimental setup (Fig 2) considerably differed depending on the lyssavirus, the route of infection, and the infectious dose (Fig 3). In the low dose i.m. groups, survival rates varied between 100 % (RABV-Raccoon, EBLV-1) and 0 % (BBLV) (Fig 3A). In contrast, in the high dose i.m. groups, all mice infected with ARAV, GBLV, IRKV, RABV-DogA and BBLV developed clinical symptoms and had to be euthanized. All animals survived after DUVV infection (Fig 3B). While following high dose inoculation with ABLV and RABV-Raccoon 67 % and 50 % of mice survived, respectively, survival rates in the remaining five groups amounted to 17 %. In contrast, when comparing bat lyssaviruses with classical RABV strains, there was no significant difference in survival rates among i.m. low dose (p=0.0862, Log-rank (Mantel-Cox) test) and high dose (p=0.8761, Log-rank (Mantel-Cox) test) infected groups. All mock-infected control mice did not develop any clinical signs and survived until the end of the observation period. All Mice inoculated i.c. with the different isolates developed clinical symptoms and were euthanized, except one mouse inoculated with BBLV, which survived until the end of the observation period (Fig S2A). The survival rate in groups inoculated i.n. with the lyssavirus isolates varied between 33 %, (EBLV-1, EBLV-2 and BBLV) and 100 %, (RABV-Racoon, RABV-Insectbat, RABV-Vampbat and IRKV, Fig S2B).

**Fig 2:**
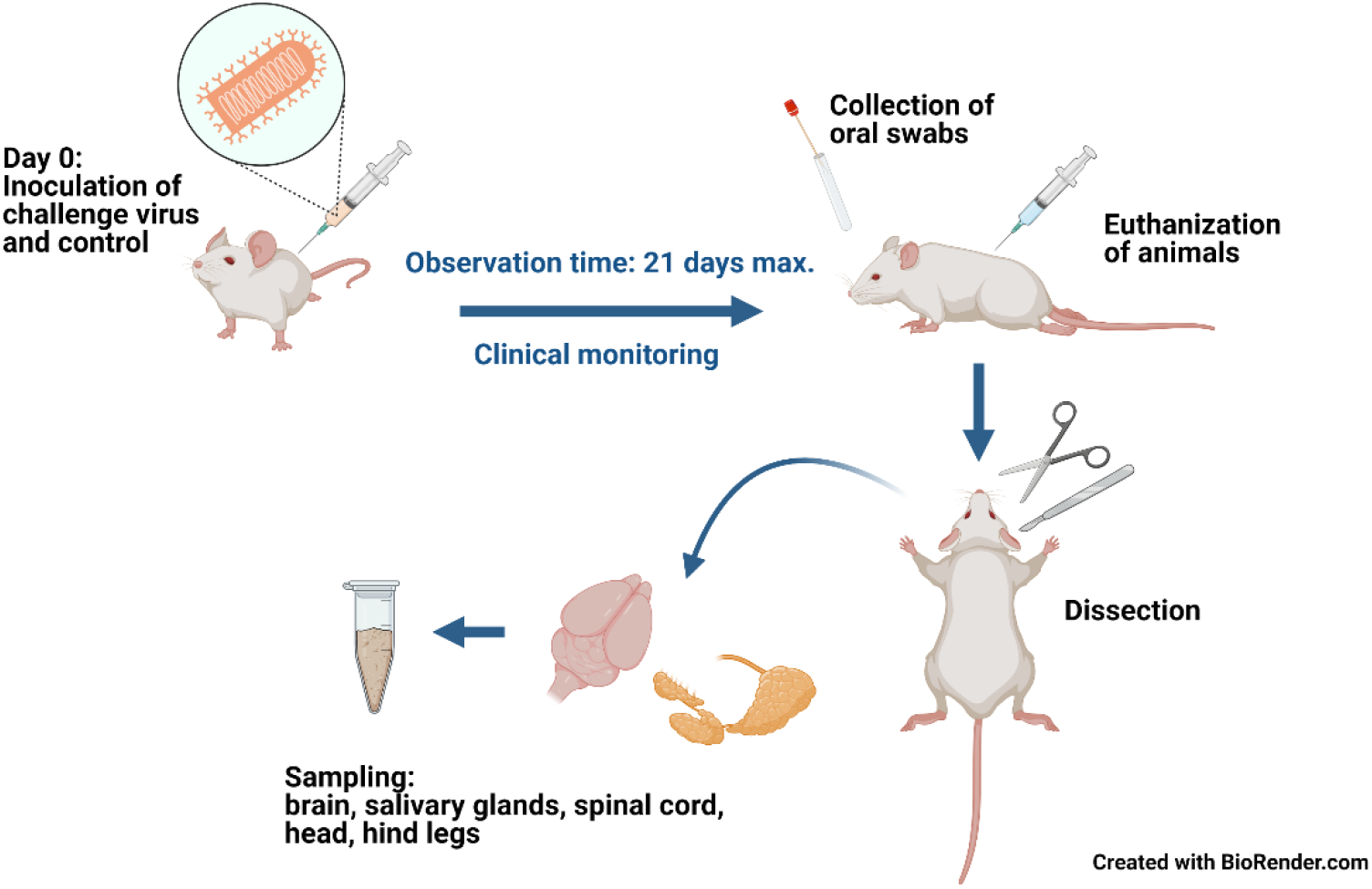
Experimental setup. Outline of the animal experiment and sample collection.

**Fig 3:**
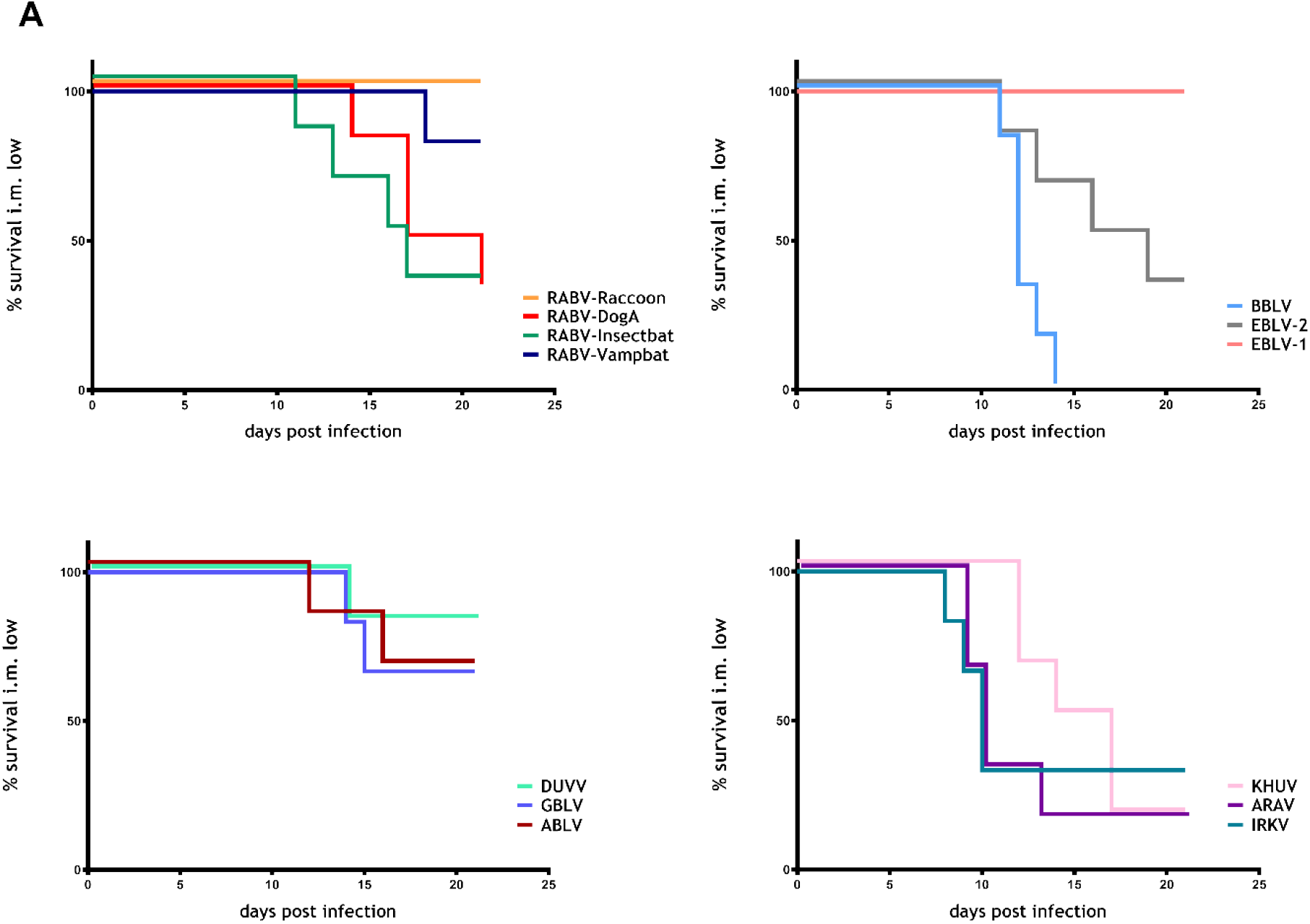
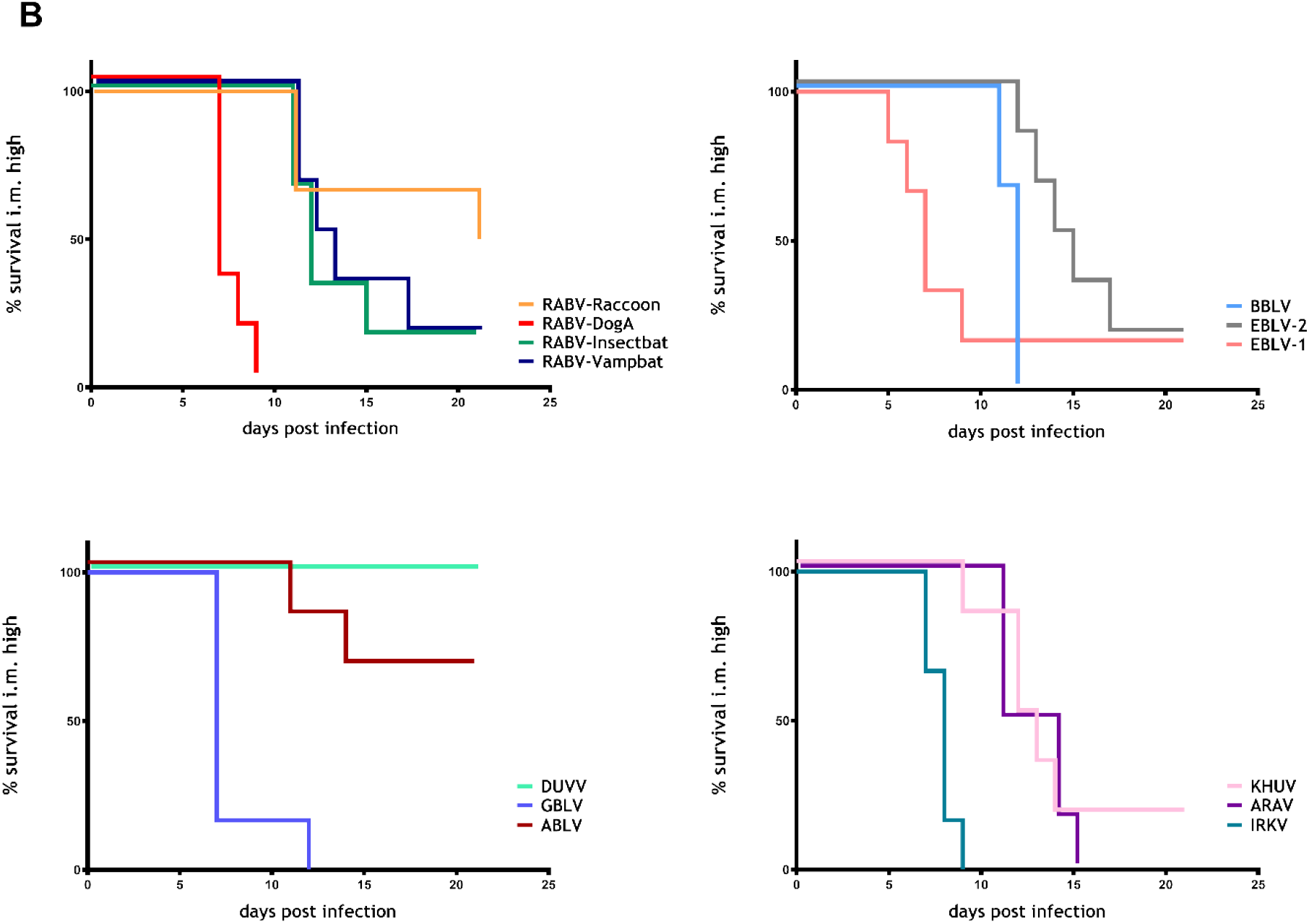
Kaplan-Meyer survival plots of the individual isolates following i.m. infection with low (A) and high doses (B). Six Balb/c mice were inoculated per group. Mock-infected control mice did not develop any clinical signs and, hence, were omitted for better visualization.

### 3.3. Incubation periods

In the i.m. low dose groups, the longest incubation periods were observed for mice inoculated with the reference strain RABV-DogA (mean: 17 days, SD±3) and with RABV-Vampbat (mean: 17 days, SD±0). However, in the group inoculated with RABV-Vampbat only a single animal developed clinical signs at all, the same applied to mice inoculated with DUVV. Incubation periods of mice inoculated with BBLV (mean: 10 days, SD±1), ARAV (mean: 9 days, SD±1.8), IRKV (mean: 8 days, SD±0.5) were significantly shorter compared to RABV-DogA (Fig 4A). In contrast, all mice infected with RABV-Raccoon and EBLV-1 survived until 21 dpi showing no clinical signs at all. Differences in incubation periods in the i.m. high dose groups were more pronounced. Mice inoculated with RABV-DogA (mean: 7 days, SD±0.8), EBLV-1 (mean: 7 days, SD±1.5), BBLV (mean: 9 days, SD±0.8), IRKV (mean: 6 days, SD±0.4) and GBLV (mean: 8 days, SD±2.3) displayed clinical signs earlier (mean <10 days) than the remaining groups (Fig 4B). Regarding classical and bat-associated RABVs, the mean incubation for RABV-DogA was significantly different to those for RABV-Raccoon (mean: 13 days, SD±5.8) and RABV-Vampbat (mean: 12 days, SD±2.8) (Fig 4B). None of the mice infected with DUVV (high dose; i.m.) showed clinical signs until the end of the observation period.

**Fig 4:**
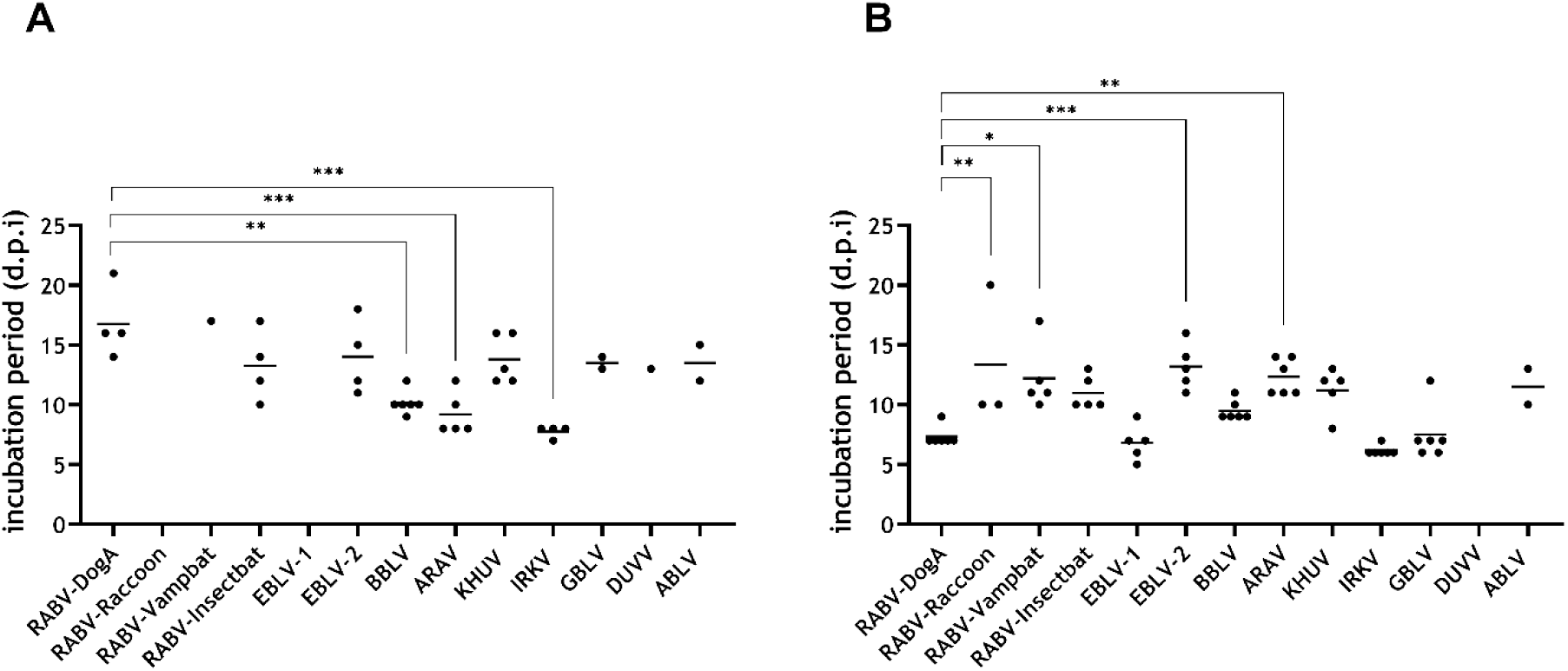
Incubation periods after low dose (A) and high dose (B) i.m. infection. Mean values are provided as horizontal lines. Statistical differences between the mean of RABV-DogA as a reference challenge strain and the means of other lyssaviruses are indicated (* p ≤ .05; ** p ≤ .01; *** p ≤. 001; ordinary one-way ANOVA with Tukey’s multiple comparison test).

### 3.4. Clinical signs

Independent of either the bat lyssavirus isolates or the classical RABV variants used, mice that succumbed to infection displayed clinical pictures suggestive of rabies. In general, clinical signs summarized as a progressive deterioration of the general health condition included decreased activity, hunched back, ruffled fur, weight loss, and lethargy or loss of alertness. Other clinically evident signs comprised paralysis, paresis, spasms, convulsive seizures, tremor, pruritus, aggressiveness, tameness, moving in circles, or extremely increased uncoordinated movements (Fig 5). For i.m. infected mice, disease progression after onset of clinical signs was either peracute with rapid development of clinical signs within <12 hours from healthy to apathetic, or disease duration was comparatively slower starting with mild, unspecific signs evolving into the full clinical picture within two to three days. The former was more frequently observed in mice inoculated with IRKV or KHUV and the latter was particularly common in mice inoculated with BBLV.

**Fig 5:**
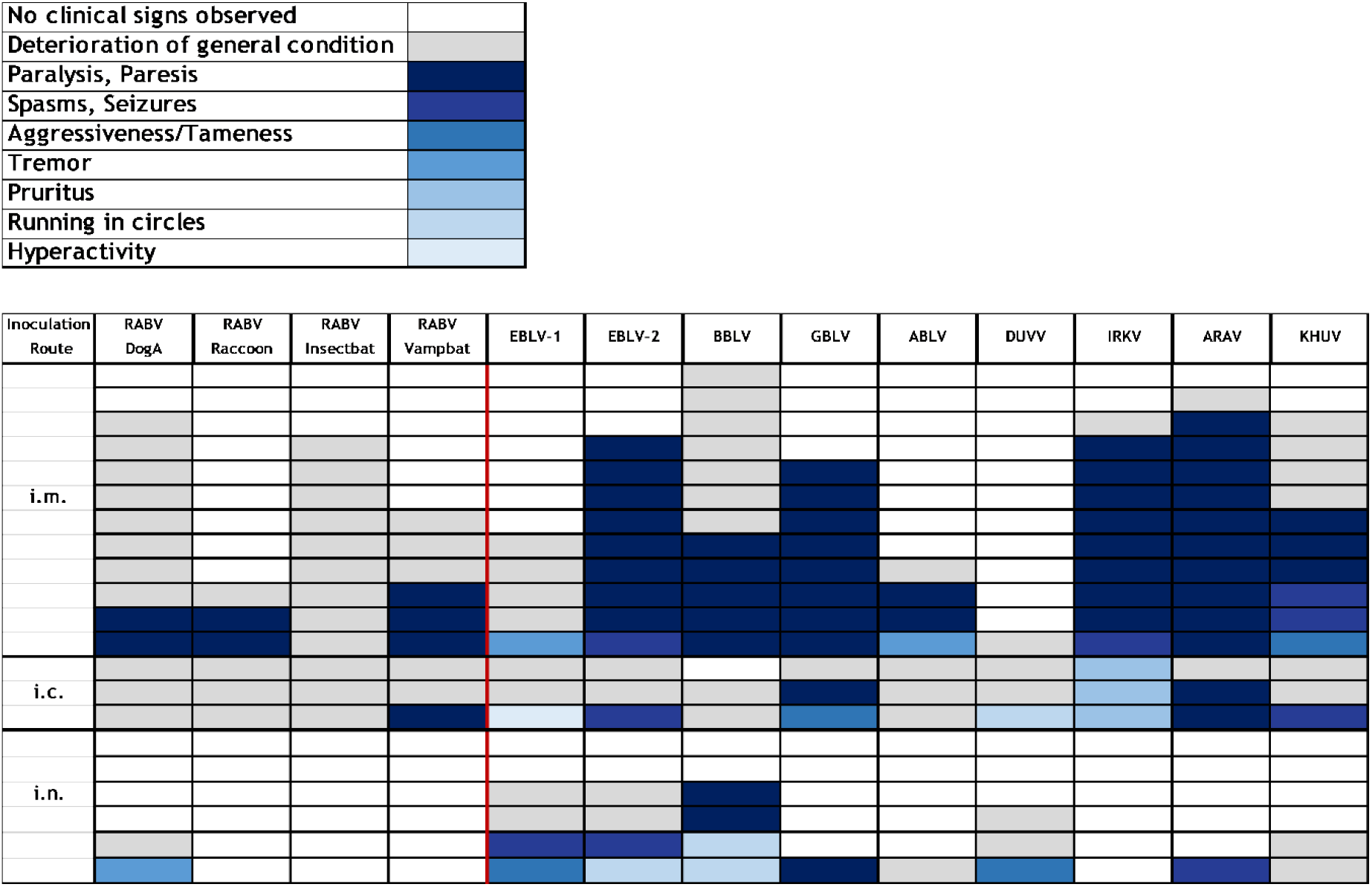
Clinical signs in individual mice inoculated with the indicated lyssaviruses by the i.m. (n= 156; 78 high dose and 78 low dose), i.c. (n= 39) or i.n. (n= 78) route of infection. The color code represents the predominant clinical sign. For clarity, high and low dose i.m. infections were combined.

After i.m. infection, 41 % of diseased mice had a deterioration of their general condition as the most prominent clinical sign, while 52 % showed paralysis and paresis. The latter was more pronounced in GBLV (100 %), ARAV (91 %), EBLV-2 (89 %) and IRKV (80 %), whereas EBLV-1 and DUVV are outliers in the non-RABV lyssaviruses with no paresis/paralysis observed in diseased mice. Generally, with 63 % over 25 %, paresis/paralysis was significantly increased in non-RABV after i.m. infection of mice (*χ*^2^=21.16, p<0.0001). Also, paresis/paralysis occurred generally less often in i.c. (11 %)- and i.n. (14 %)-infected mice (Fig 5).

### 3.5. Index-based comparative pathogenicity

To allow for a ranking in pathogenicity, we implemented a novel intramuscular pathogenicity index (IMPI), which takes clinical signs and deaths/euthanasia of all i.m.-inoculated (dose-independently) mice into account. If all infected mice died at day 1, the IMPI would be 2, whereas it would be 0 if all mice survived with no clinical score. Depending on the lyssavirus used, indices ranging between 1.14 and 0.07 were obtained (Fig 6). The IMPI scored highest for IRKV and BBLV (>1) compared to the classical RABV-DogA (0.85). In contrast, RABV-Raccoon and DUVV had the lowest score (<0.19) (Fig 6).

**Figure 6:**
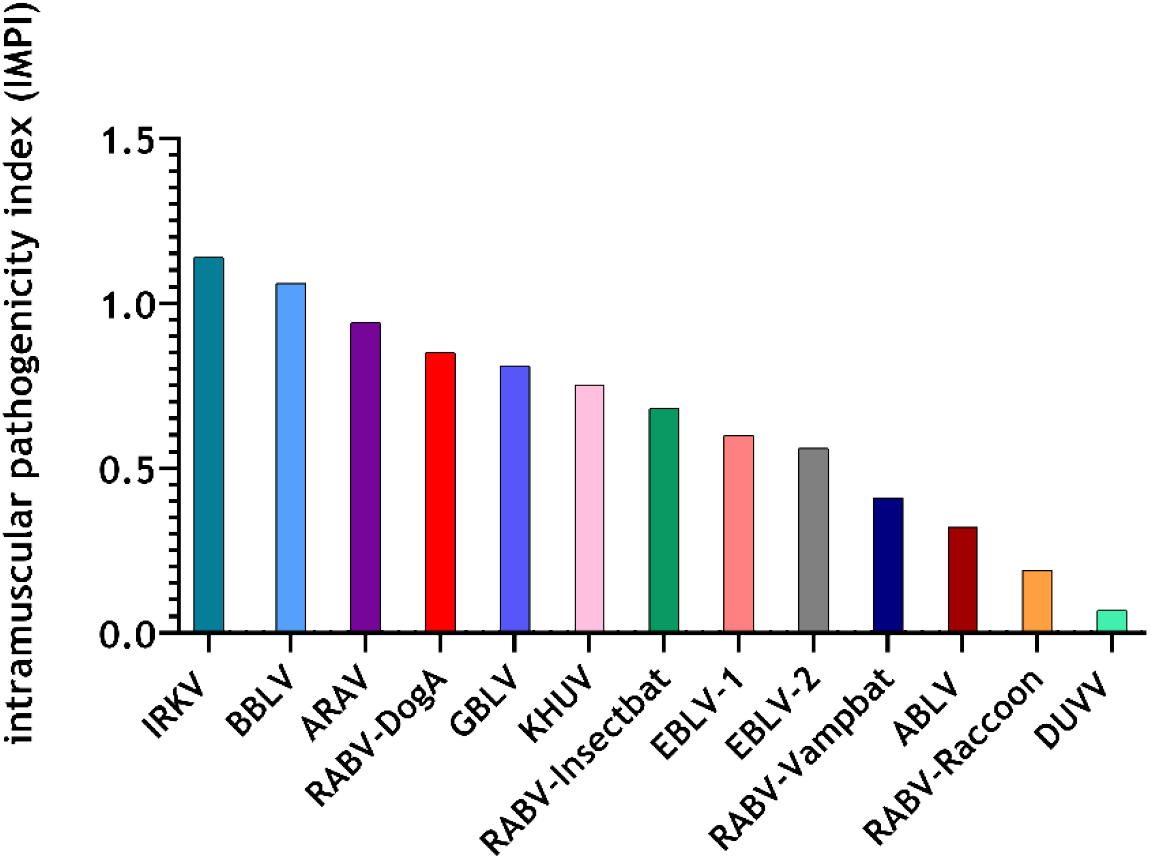
Intramuscular pathogenicity index (IMPI) of the different lyssavirus isolates obtained in the mouse model. Depicted are pathogenicity indices of combined datasets of i.m. low and high dose infected animals. A maximum index value of 2 would be reached if all mice died at day 1 post-infection.

### 3.6. Infection of neuron and astrocytes by selected bat lyssaviruses

Since pathogenic RABV have recently been shown to exhibit a specific astroglia tropism [42], we analyzed whether other lyssaviruses could infect central nervous system resident astrocytes to a comparable extent and whether differences in the pathogenicity correlate with astroglia infection levels. Therefore, the brain cell tropism of selected bat lyssaviruses with a high (IRKV, BBLV) and a low (RABV-Vampbat, DUVV) pathogenicity index (Fig 6) was investigated by 3D immunofluorescence imaging and quantitative analysis of infected neurons and astrocytes (Fig 7).

**Fig 7:**
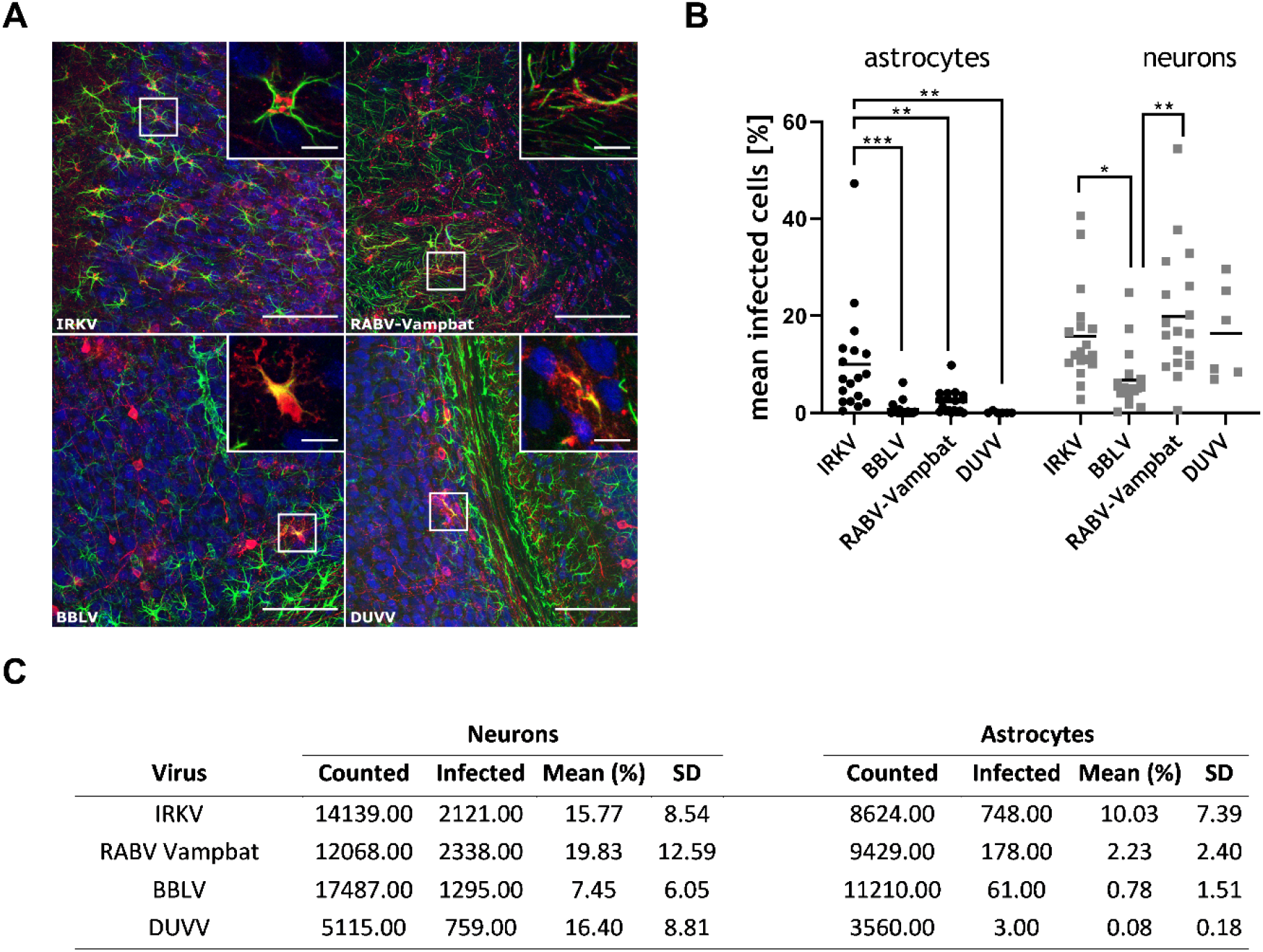
Comparison of the astrocyte tropism of different bat lyssaviruses with a high and low IMPI. **A)** Indirect immunofluorescence of solvent-cleared brain sections demonstrates the presence of lyssavirus phosphoprotein P (red), neurons (blue, marker: NeuN) and astrocytes (green, marker: glial fibrillary acidic protein, GFAP). Insets show RABV P accumulation (red) at GFAP-positive cells (green). x, y = 387.5 µm, 387.5 µm; z = 65.5 µm (IRKV), 36.5 µm (RABV-Vampbat), 64.5 µm (BBLV), 65.5 µm (DUVV). Scale bar = 100 µm (overview), 15 µm (inset). **B)** Percentage of infected astrocytes (black dots) and neurons (gray squares). Per virus, 3 to 11 × 10^3^ astrocytes and 5 to 17 × 10^3^ neurons were counted in independent confocal z-stacks in two animals per isolate (one animal for DUVV). Each dot represents the ratio of infected astrocytes or neurons in an analyzed z-stack. Mean values are provided as horizontal lines. Statistical comparisons of the means between the different groups are indicated for those with a statistically significant difference (* p ≤ .05; ** p ≤ .01; *** p ≤. 001; ordinary one-way ANOVA with Tukey’s multiple comparison test). **C)** Corresponding data table for the measurements.

Intramuscular infections with IRKV, RABV-Vampbat, BBLV and DUVV resulted in a mean of 15.77 % (SD±8.54), 19.83 % (SD±12.59), 7.45 % (SD±6.05), and 16.4 % (SD±8.81) virus-positive neurons, respectively, demonstrating that IRKV and RABV-Vampbat had significantly higher levels of neuron infection compared to BBLV in clinically diseased mice (Fig 7B, C). Concerning astroglia infection, IRKV-infected mice featured the highest percentage of infected astrocytes (10.03 %; SD±7.39). While astrocyte infection was lower in RABV-Vampbat (2.23 %; SD±2.4)- and BBLV (0.78 %; SD±1.51)-infected mice, only individual infected cells were identified in the DUVV-infected mice (Fig 7B, C).

### 3.7. Virus shedding – detection of viral RNA and viable virus in salivary glands and oral swabs

The detection of viral RNA in salivary glands and oral swabs in mice differed depending on the lyssavirus species used for inoculation. Positivity rates for all salivary gland and oral swab samples were highest for RABV-DogA (100 %), followed by RABV-Insectbat (92 %), RABV-Raccoon (33 %) and RABV-Vampbat (33 %) (Fig 8A). Independent of the route of infection, 97 % of diseased mice inoculated with RABV strains were positive for viral RNA in salivary glands, and in 72 % of mice viral RNA was also detected in the corresponding oral swabs (Fig 8B). In contrast, mice infected with non-RABV bat lyssaviruses exhibited significantly lower positivity rates (p<0.0001, Fischer’s exact test), i.e. in only 50 % of the diseased animals, salivary glands were positive for viral RNA, and 12 % exhibited both virus RNA-positive salivary glands and oral swabs (Fig 8B). Regarding the presence of infectious virus, a similar pattern was observed when all routes of infection were considered (Fig 8C, D). When grouped together, infectious virus could be isolated in 86 % of salivary gland samples from mice that succumbed to RABV, while infectious virus shedding, as determined by virus isolation from both salivary glands and oral swabs, was 47 % overall, with the highest proportion in RABV-DogA (92 %) and RABV-Insectbat (67 %). While BBLV- and GBLV-diseased mice had positive salivary glands at frequencies of 83 % and 67 %, respectively (Fig 8C), no virus was isolated from oral swabs. None of the salivary glands or oral swabs from ARAV-, IRKV-, and KHUV-infected mice were positive for viral RNA or infectious virus (Fig 8A, C). Combined, similar to the presence of viral RNA, mice infected with non-RABV bat lyssaviruses exhibited significantly lower (p<0.0001, Fischer’s exact test) positivity rates for infectious virus isolation as compared to RABV-infected mice (Fig 7D).

**Fig 8:**
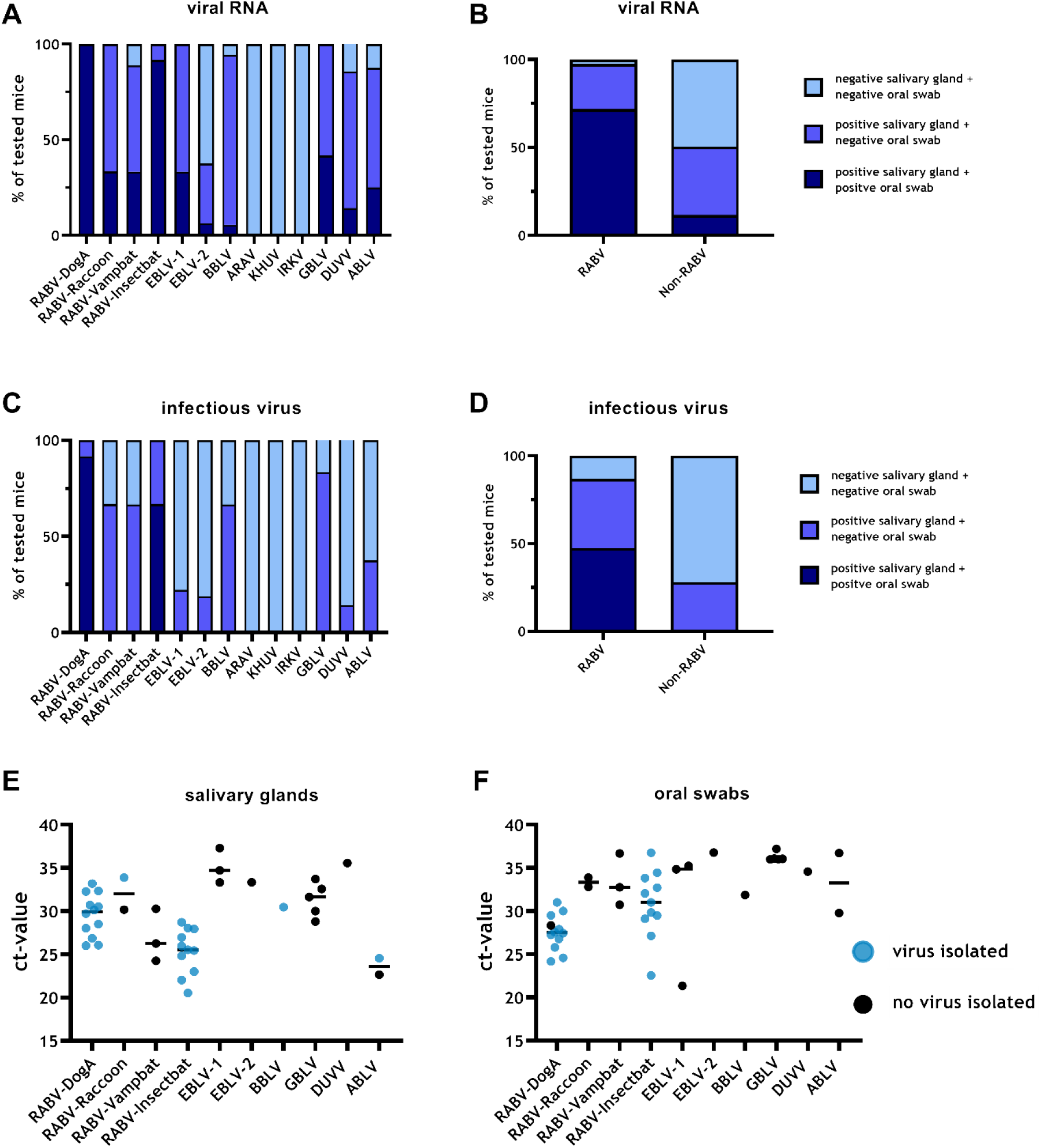
Comparison of virus shedding in lyssavirus-infected mice. Percentage of animals positive/negative for viral RNA (A, B) and viable virus (C, D) in either salivary glands or oral swabs or both according to individual lyssaviruses (A, C) or grouped according to RABVs and non-RABV bat lyssaviruses (B, D). **Correlation between ct-values as obtained in RT-qPCR and results of virus isolation in salivary glands (E) and oral swabs (F)**. Here, only animals were considered were active shedding (positive salivary gland and positive corresponding oral swab) was observed. Individual ct-values are shown and the mean is indicated. Successful virus isolations in cell culture are highlighted.

When data were analyzed according to inoculation dose and route, shedding of infectious virus was observed more often in mice diseased after i.c. inoculation (51 %), followed by i.m. high dose (41 %), i.n. (35 %), and i.m. low dose (29 %). The mean ct-values were significantly lower in samples with successful virus isolation from salivary glands (p=0.003, unpaired t-test, Fig 8E) and corresponding oral swabs (p= 0.003, unpaired t-test, Fig 8F) as opposed to unsuccessful virus isolation. This observation is mainly driven by the low ct-values observed in RABV-Insectbat and RABV-DogA.

## 4. Discussion

Although lyssaviruses comprise a genetically close group of viruses, which all cause the disease rabies, differences in their phenotype may indicate different risks for veterinary and public health. In our assessment, 13 different lyssaviruses exhibited differences in their replication kinetics in terms of the growth dynamic and the final virus titer. As such, in vivo replication kinetics may reflect replication in the animal host and thus may explain differences seen in incubation periods and pathogenicity among individual lyssaviruses. Interestingly, relatively short incubation periods (IRKV, EBLV-1) (Fig 4A, B) or relatively high (IRKV, BBLV) or low pathogenicity indices (ABLV) (Fig 6) in the animal model also correlate with the replication kinetics (Fig 1).

Similarly, the number of animals that survived infection with the individual lyssaviruses varied (Fig 3A, B). Survival was not associated with belonging to classical RABV as opposed to non-RABV bat lyssaviruses. There was considerable variation seen in incubation periods across lyssavirus species as well as between animals of individual groups (Fig 4A, B). However, we have no evidence that bat-associated lyssaviruses cause longer incubation periods in mice compared to observations made in epidemiological bat models [59,60], and supporting case studies [61,62].

The perception and subsequent recording of clinical signs might be biased due to the clinical score being applied for animal welfare reasons and due to the temporally restricted observation scheme. However, all investigated lyssaviruses caused a clinical picture and eventually death in mice, albeit at different scale. We observed the clinical outcome to be predominantly dependent on the virus species but also on the route of application. The fact that clinical signs such as paralysis and spasms were more pronounced after i.m. inoculation corroborates previous findings [34]. On the other hand, the observation that mice infected i.m. with non-RABV bat lyssaviruses were more likely to develop spasms and paralysis is interesting but requires further investigation. While lyssavirus species-dependent clinical signs were reported before [36], in summary, no clear pattern in regard to particular lyssavirus species was evident in our study (Fig 5).

Overall, the variation in pathogenicity factors shown in our study highlights the complexity and difficulty to establish a holistic concept for the classification of lyssaviruses. To integrate these factors from our standardized in vivo model, we applied an intramuscular pathogenicity index for lyssaviruses, and thereby demonstrated a high diversity across phylogroup I lyssaviruses that had not been shown to such an extent before. Remarkably, non-RABV bat lyssavirus isolates are among those with the highest IMPI (Fig 6), questioning previous suggestions that bat-related lyssaviruses are less pathogenic [63]. Interestingly, IRKV demonstrated the highest pathogenicity in a ferret model as well in comparison to the other bat-associated lyssaviruses KHUV and ARAV [13]. Moreover, the fact that current RABV-based biologicals provide only partial protection against IRKV challenge [13,64] further emphasizes the higher risk for a lethal outcome associated to IRKV infections. Even though EBLV-1, DUVV, and IRKV have all caused human rabies cases [10,65], no elevated pathogenicity was found in our model using isolates from human cases (EBLV1, DUVV) (Fig 6). Of note, the Yuli isolate we used here is the only available EBLV-1 isolated from a fatal human case [66].

Using high-resolution confocal laser-scanning microscopy, the CNS cell tropism for bat-related lyssaviruses was analyzed here for the first time and provides new insights into their capability to infect astrocytes (Fig 7). A comparison between RABV field isolates and lab-adapted strains revealed that astrocyte infection after i.m. inoculation is associated with field strains and, thus, might be a potential pathogenicity determinant [42]. In our analyses using representatives of bat-associated lyssaviruses, mice infected with IRKV, as a representative for high pathogenicity (IMPI=1.14), had significantly higher proportions of infected astrocytes than RABV-Vampbat (IMPI=0.41)- and BBLV (IMPI=1.06)-infected mice. Interestingly, only sporadic astrocyte infection was found in DUVV (IMPI=0.07)-infected mice. Even though the restricted number of lyssaviruses analyzed for astrocyte infection hinders a full comparison, the results support previous studies of RABV on the association of astrocyte tropism and pathogenicity. Additional virus isolates and strains have to be analyzed in further studies to confirm the role of astrocyte tropism in lyssavirus pathogenicity. However, different virus kinetics and astrocyte-related innate immune reactions may affect the progression kinetics, immune pathogenicity, and further spread of the virus to peripheral salivary glands. The latter may represent a key issue in terms of virus transmission and maintenance in host populations. In our analyses, virus shedding was not demonstrated in IRKV-infected mice. In general, virus shedding represents a striking discrepancy between RABV and other bat lyssaviruses, as virus shedding was significantly reduced in non-RABV bat lyssaviruses in the mouse model. Interestingly, while shedding was highest for the dog rabies strain RABV-DogA, bat-related RABV isolates and a raccoon RABV variant demonstrated a lower percentage of active shedding (Fig 7A-D). Of note, the raccoon RABV lineage in the Americas is a result from an ancient sustained spillover event from bats [67].

By our definition, active shedding is the successful virus isolation from salivary glands and the respective oral swab sample. Of note, a positive result from an oral swab may not necessarily correspond to viral excretion in saliva itself because the swab could also contain desquamated cells, including infected neurons, which may not necessarily have been excreted naturally. However, technically it was not possible to extract only saliva and the same methodology was used for all isolates. Discrepant results in virus isolation in salivary gland samples and corresponding oral swabs can be explained by the fact that neurons innervating the glands are infected, without excreting virus in the lumen of the gland. The reason for the limited shedding of bat-associated lyssaviruses may be a yet unknown block or barrier in virus distribution in the salivary gland of non-bat mammals. Another explanation could be intermittent shedding, i.e. samples might have been taken at time points when virus was temporarily not shed, but the salivary gland was still virus positive. Nevertheless, intermittent shedding does also apply for RABV strains and consequently it does not completely explain the differences. For some isolates the disease duration in mice was very short so that the centripetal spread of virus may not have reached the salivary gland before death or euthanasia. Interestingly, neither viral RNA nor infectious virus could not be found in salivary glands or oral swabs for the three virus species IRKV, KHUV, and ARAV. Furthermore, there was a gradient for virus shedding with lowest percentage of shedding in low dose i.m. infected animals, and highest in i.c. inoculated ones. While the mean ct-value in successfully isolated samples was significantly lower, there was a high range in ct-values between 20 and 37 (Fig 8E, F), and no threshold for successful isolation could be defined.

In any case, our shedding results could mimic the capability of transmission from a spillover host to another conspecific as the prerequisite of a sustained spillover [41], and thereby contribute to an overall risk assessment for lyssaviruses. Our shedding results in the mouse model confirm previous assumptions based on field observations that the potential for sustained spillovers is highest in RABV as opposed to other bat lyssaviruses [8].

Animal models have greatly improved our understanding of virus-host interactions. When studying virus-host interactions of bat-related lyssaviruses, experimental studies in bats as their primary reservoir hosts would be ideal. While the results of those studies can infer the pathobiology in the host, they are particularly challenging due to their demanding housing conditions, difficult handling, conservation issues and partly strict protection status [68]. Furthermore, they are less suitable to assess the pathogenicity in non-reservoir hosts. Therefore, for reasons of comparison, the selection of reliable and standardized alternative animal models is of great importance in the study of pathogenicity factors. While mice are considered a standard model for studying lyssavirus pathogenicity [69], in fact little attention has been paid to comparability. Here, our model facilitates a direct comparison of pathogenicity data of 13 different phylogroup I bat lyssaviruses by using a consistent approach regarding experimental conditions, e.g. cells for viral propagation, mouse breed, viral doses, inoculation routes as well as observation times and scoring schemes. We thereby optimized an in vivo mouse model established for EBLV-1 [34] by including data on virus shedding. Furthermore, we established a novel matrix for comparing pathogenicity based on clinical parameters, the intramuscular pathogenicity index IMPI. Such indices are commonly used for avian influenza viruses and Newcastle disease viruses to directly infer the potential for causing disease in animals and thus relate to the respective veterinary control measures. In our case for lyssaviruses, the index can also be used to summarize the pathogenic potential of the individual lyssavirus isolate.

We included i.n. and i.c. inoculation routes in our assessment (Fig S2), with i.c. primarily used as a positive infection control with the low dose virus inoculum, whereas i.n. application was tested as it was speculated before that virus transmission via aerosols could contribute to disease spread among bats and in spillover infection [68,70]. While i.n. application led to infection of some animals (S2B Fig) likely via the olfactory pathway [71], no clear indication for a specific role in bat lyssaviruses was found (Fig 5), supporting experimental studies in bats [72,73]. Therefore, we then focused our analyses on pathogenicity to i.m. inoculation as the most likely route of virus infection. For virus shedding, we wanted to assess the virus’ potential to be transmitted when an animal develops disease, and therefore all inoculation routes were considered. Nevertheless, there are some limitations in our study. The number of mice used was kept to a minimum, respecting animal welfare guidelines on the 3R principle [74]. Also, propagation of viruses in cell culture was a necessary requirement to obtain sufficient virus stocks for the experiments. To minimize the possibility of adaption to cell culture that may influence the results obtained, we used Na 42/13 cells, a mouse neuroblastoma cell line considered to be primary target cells for lyssaviruses, and kept the number of passages as low as possible. Also, the likelihood of adaptive mutation is regarded low since lyssaviruses, i.e. RABV [75] and EBLV-1 [76] have among the lowest mutation rates of RNA viruses [77]. The full genome sequences derived from passaged material did not give evidence of nucleotide exchanges compared to other sequences of the primary isolates.

## 5. Conclusion

Here, we have comparatively assessed the pathogenicity and virulence of a wide diversity of lyssaviruses belonging to phylogroup I using a standardized and systematic approach. Interestingly, we found in our model bat-associated lyssaviruses, which are more pathogenic and virulent than a classical RABV challenge strain. In fact, no tendency towards a generally reduced pathogenicity of bat-associated lyssaviruses as opposed to classical RABV can be confirmed and thus each isolate should be considered individually concerning its pathogenicity. In contrast to RABV, we could not determine virus shedding in all other lyssavirus-infected mice. This indicates a limited potential of those lyssaviruses to spread beyond the initial spillover-host, and may explain the absence of onward transmission in non-RABV bat lyssaviruses.

## Acknowledgements

The technical assistance by Jeannette Kliemt and Patrick Zitzow is gratefully acknowledged. Also, we would like to thank all animal keepers at FLI for supporting the experimental studies.

## Notes

### Competing Interest Statement

The authors have declared no competing interest.

